# Triacylglycerol synthesis enhances macrophage inflammatory function

**DOI:** 10.1101/2020.02.03.932079

**Authors:** Angela Castoldi, Lauar B Monteiro, Nikki van Teijlingen Bakker, David E Sanin, Nisha Rana, Mauro Corrado, Alanna M Cameron, Fabian Hässler, Mai Matsushita, George Caputa, Ramon I. Klein Geltink, Jörg Büscher, Joy Edwards-Hicks, Erika L Pearce, Edward J Pearce

**Affiliations:** Department of Immunometabolism, Max Planck Institute of Epigenetics and Immunobiology, Freiburg im Breisgau, Germany; Faculty of Biology, University of Freiburg, Freiburg im Breisgau, Germany; Metabolomics Facility, Max Planck Institute of Epigenetics and Immunobiology, Freiburg im Breisgau, Germany

**Keywords:** Lipid droplets, macrophages, triacylglycerol, acylcarnitine, PGE2, inflammation

## Abstract

Macrophages are integral to most tissues. Foam cells, macrophages with lipid droplets (LDs) which are stores of triacylglycerols (TGs) and cholesterol esters (CEs), are found in various disease states^1^. LDs can act as energy stores since TG lipolysis releases fatty acids (FAs) for mitochondrial oxidation (FAO), a process that relies on long-chain FA conversion into acylcarnitines by the enzyme Cpt1a^2^. However, in macrophages, proinflammatory signals result in diminished FAO and increased TG synthesis with LD development^3,4^. We explored the significance of LDs in cells that do not utilize FAO. We show that macrophages stimulated with lipopolysaccharide (LPS) plus interferon-γ (IFNγ) accumulate TGs in LDs, and long-chain acylcarnitines. In these cells, inhibition of TG synthesis results in diminished LD development, and increased long chain acylcarnitine levels, suggesting that FA fate is balanced between TG and acylcarnitine synthesis. Nevertheless, TG-synthesis is required for inflammatory macrophage function, since its inhibition negatively affects production of proinflammatory IL-1β, IL-6 and PGE2, and phagocytic capacity, and protects against LPS-induced shock in vivo. Failure to make PGE2 is critical for this phenotype, since exogenous PGE2 reverses the anti-inflammatory effects of TG-synthesis inhibition. These findings place LDs in a position of central functional importance in inflammatory macrophages.

---

Inflammation has critical protective functions, but when unregulated can also cause disease, in which IL-1β and other proinflammatory cytokines are implicated^5^. Recent work has established a strong link between inflammatory activation signals and induced changes in macrophage metabolism that are essential for the cells to perform their subsequent functions^6^. Major components of this metabolic reprogramming include enhanced Warburg metabolism, associated with diminished oxidative phosphorylation (OXPHOS) and altered TCA cycle dynamics in which glucose carbon is redistributed via citrate/aconitate to the synthesis of fatty acids (FA) and itaconate^7^. Increased FA synthesis in inflammatory macrophages is associated with additional broad changes in lipid metabolism^8^, including the accumulation of TGs and CEs^3,9^, which are stored within LDs. TG synthesis depends on the acyl-CoA:diacylglycerol acyltransferases 1 and/or 2 (DGAT1,2)^10^, which catalyze the covalent addition of a fatty acyl chain to diacylglycerol (DG)^10^. LDs are the core energy storage organelles of adipocytes, but develop in other cell types as well, where they can again act as energy stores for fueling cell intrinsic ATP production via mitochondrial FAO^2,11^. However, LDs are recognized to mediate additional functions including the sequestration of toxic lipids and the prevention of excessive endoplasmic reticulum (ER) stress, and as lipid donors for autophagosome formation^2,12,13^.

*In vivo*, LD-containing macrophages are most well recognized in atherosclerotic lesions and in tuberculosis^1,14^ More recent work has highlighted the presence of foam cells in active multiple sclerosis, certain cancers, white adipose tissue during obesity, and in bronchoalveolar lavage from individuals suffering from vaping-related lung disease^1^. The ratio of TGs to CEs in LDs in different settings is likely to be important, and there is ongoing discussion regarding whether macrophages that contain CE-rich LDs are universally proinflammatory^15,16^.

Inflammatory macrophages increase the commitment of resources to FA synthesis^7^. Consistent with this, these cells accumulate LDs^3,9,4^ but the function of LDs in this setting is unclear, especially since inflammatory macrophages do not use FAO^3,4^.We began to address this by investigating the process and functional significance of TG synthesis in response to the strong proinflammatory stimulus provided by LPS plus IFNγ (Fig. 1a). We found that the expression of both *Dgat1* and *Dgat2*, the enzymes responsible for synthesizing TGs from DGs, increased in inflammatory macrophages as part of a broader transcriptional signature of genes involved in TG synthesis and utilization (Fig. 1a), but that *Dgat1* was most strongly expressed (Fig. S1a). As expected, TG and LD accumulation were also marks of inflammatory activation (Figs. 1b, c, d). Consistent with their use for TG synthesis (Fig. 1a), overall levels of free FAs and DGs were diminished in inflammatory compared to resting macrophages (Fig. 1e). Lipidomics revealed broad, dynamic changes in lipids as a result of inflammatory activation, with the accumulation of phosphatidylcholines (PCs), phosphatidylinositols (PIs), phosphatidylehtanolamines (PEs), lysophosphotidylcholines (LPCs), sphingomyelins (SMs), CEs and hexosylceramides (HEXCERs) beginning as early as 2 h post activation (Fig. S1 b). This was not the case in IL-4-stimulated alternatively activated macrophages (Fig. S1b), which actively fuel FAO by lipolysis and which, unlike inflammatory macrophages, do not contain visible LDs^4^.

**Figure 1.**
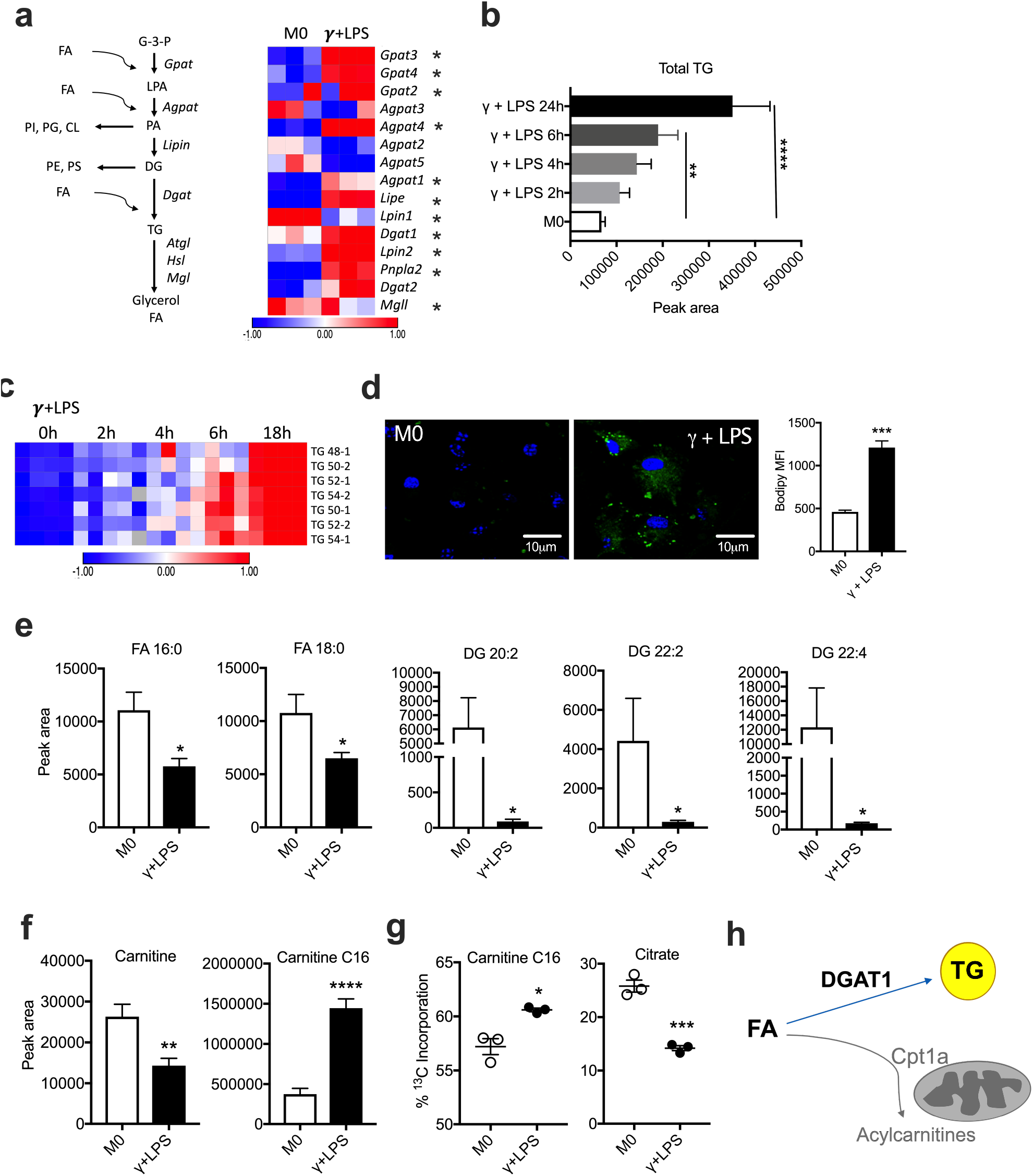
Inflammatory activation results in the remodeling of fatty acid metabolism. **(a)** TG pathway and expression of pathway genes, as measured using RNAseq, in resting (M0) and inflammatory macrophages (stimulated with IFNγ + LPS for 18 h). (b, c) Time-dependent accumulation of total TG **(b)**, and different TG species **(c)**, in macrophages after IFNγ + LPS stimulation, measured using mass spectrometry (MS). **(d)** Bodipy 493/503 staining in resting (M0) macrophages and macrophages stimulated with IFNγ + LPS, for 18 h, visualized by confocal microscopy and measured by flow cytometry (MFI, mean fluorescence intensity). **(e)** Fatty acids (FA), and diacylglycerols (DG), **(f)** carnitine and C16 acylcarnitine measured using MS, and **(g)** total fraction contribution (FC) of palmitate carbons into C16 acylcarnitine and citrate. Resting (M0) macrophages and macrophages stimulated with IFNγ + LPS were cultured in the presence of 100μM of ^13^C-palmitate for 18h, measured using MS. **(h)** Inflammatory macrophages convert FA to TG which are stored in LDs (blue arrow), or to acylcarnitines through Cpt1a in the outer mitochondrial membrane (gray arrow). This manuscript is focusing on the significance of TGs for inflammatory macrophage activation. Error bars are mean ± SEM (*p < 0.05, **p < 0.01, ***p < 0.001 and ****p < 0.0001 compared with control or untreated, or as indicated). (a-c), (e-g) representative of two experiments and (d) representative of tree experiments.

Despite the fact that FAO and OXPHOS are inhibited during inflammatory activation^3,17^, expression of *Cpt1a*, which encodes carnitine palmitoyltransferase 1a (Cpt1a), was increased in inflammatory macrophages (Fig. S1c). Cpt1a is an enzyme located in the outer mitochondrial membrane that converts long chain FA into acylcarnitines that can then be transported into the mitochondrial intermembrane space prior to subsequent transfer into the matrix for entry into FAO. We found that increased Cpt1a was associated with a significant drop in free carnitine correlated with the accumulation of long chain acylcarnitines (Figs. 1f, S1d). This was despite parallel increases in expression of *Cpt2*, which encodes the enzyme that catalyzes the removal of carnitine from aceylcarnitines (Fig. S1c). These changes did not correlate with utilization of FA in FAO, since while incorporation of ^13^C from labeled palmitate into C16 acylcarnitine was increased in inflammatory vs. resting macrophages, its incorporation into the TCA cycle intermediate citrate was diminished (Fig. 1g). This is consistent with previous findings that FAO declines in inflammatory macrophages. Furthermore, in inflammatory compared to resting macrophages ^13^C from ^13^C-labelled glycerol was preferentially incorporated into the TG precursor glycerol phosphate, rather than being used in glycolysis, where it is measurable as labeled 3-phosphoglycerate (Fig. S1e). Taken together, our data support the view that inflammatory macrophages divert available resources towards the conversion of FAs to TGs which are stored in LDs, or to the synthesis of long chain acylcarnitines (Fig. 1h).

To ascertain the role of TGs in inflammatory macrophages, we used T863, a selective DGAT1 inhibitor (DGAT1i)^18^ to suppress TG accumulation (Figs. 2a, b; S2a). This was accompanied by diminished LD accumulation (Fig. 2c), correlated with reduced staining with bodipy 493/503 (Fig. 2d), and diminished C16 bodipy uptake (Fig. 2e). DGATi also resulted in reduced levels of CEs and cardiolipins (CLs), and certain species of HEXCERs, PCs, PSs and SMs, although it had no effect on PIs and caused increases in LPCs (Fig. S2a). DGAT1i also caused the accumulation of substrate DGs in inflammatory macrophages (Figs. 2f; S2b), but free FA levels were unaffected (Fig. S2c), presumably because these were still being used to synthesize DGs. Suppression of DGAT1 using shRNA (Fig. S2d) resulted in effects that were similar to those mediated by DGAT1i, with diminished TG accumulation (Fig. 2g) and staining with bodipy 493/503 (Fig. 2h). We next queried the effects of loss of DGAT1 function on inflammatory activation. We found that the failure to make TGs and thereby accumulate LDs in DGAT1i-treated or Dgat1-shRNA-transduced LPS/IFNγ-stimulated cells was accompanied by a significant reduction in inflammatory capacity, as measured by IL-1γ (Figs. 2i, j; S2e) and IL-6 production (Figs. 2k, l; S2f). *Il1b* and *Il6* mRNAs were also diminished by DGAT1i (Fig. S2g). These data are consistent with a previous report that DGATi diminished inflammatory cytokine production in an *M. tuberculosis-infected* human macrophage cell line^19^. Additionally, we observed marked reductions in mRNAs encoding the Macrophage Inflammatory Proteins 1α (*Ccl3*) and β (*Ccl4*), the inflammasome Nalp1 (*Nlrp1b*), and reductions in expression of additional genes involved in the inflammatory response including *Ido1* and *Il1a* in inflammatory macrophages as a result of DGAT1 inhibition (Fig. S2g), attesting to the proinflammatory effects of TG synthesis. The reduction in *Il1a* expression is of interest since IL-1α was strongly implicated in the pathophysiology of atherosclerosis associated with foamy macrophage development in oleate-fed mice^20^. Earlier work showed that interfering with LD dynamics through the inhibition of TG hydrolysis negatively impacted the ability of macrophages to engage in phagocytosis^21^. Here, we observed reduced phagocytic ability following loss of DGAT1 function (Figs. 2m, S2h).

**Figure 2.**
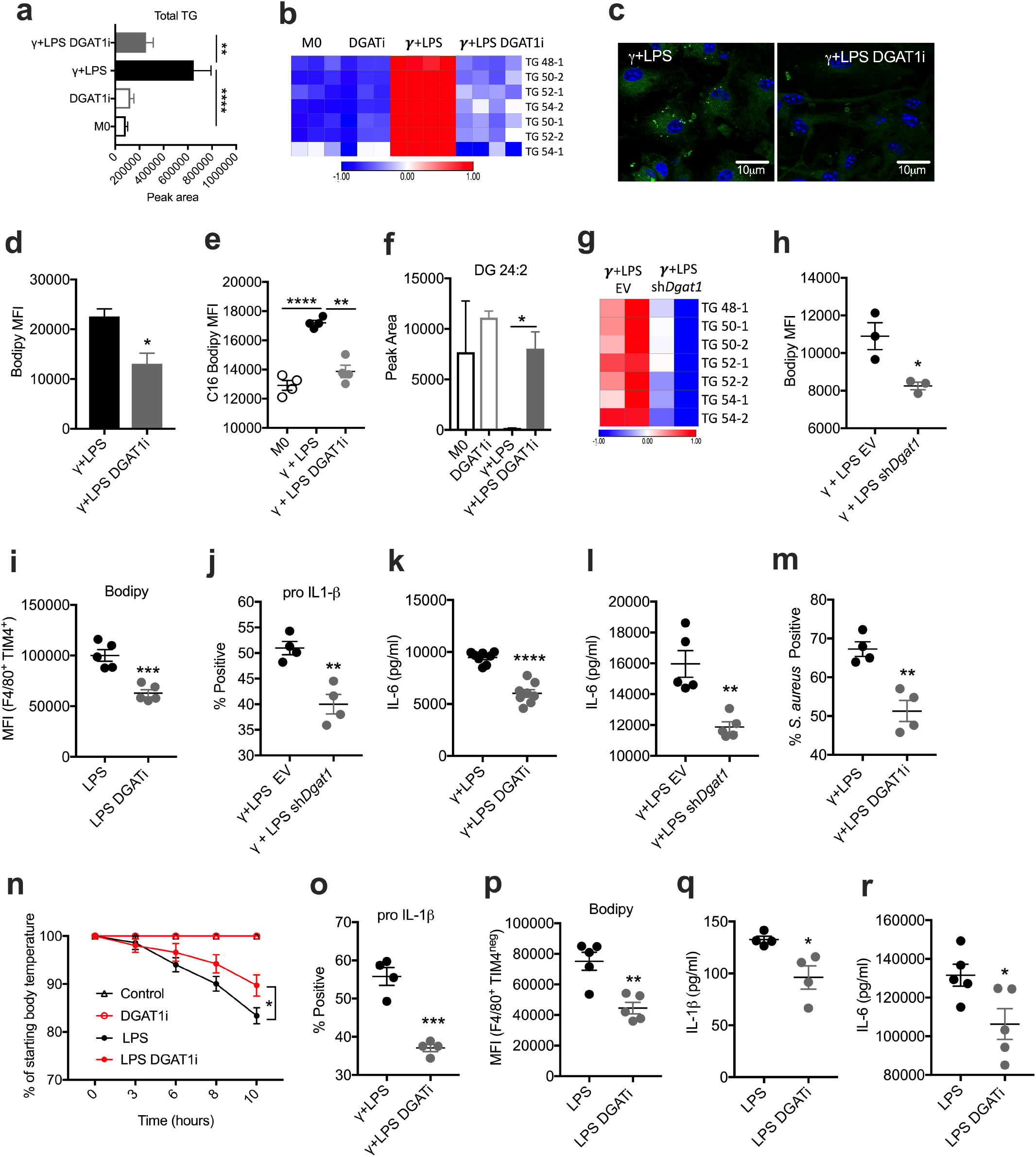
Loss of DGAT1 function decreases TG accumulation, LDs and inflammatory function. **(a, b)** Effect of DGAT1 inhibitor T863 (DGAT1i) on total TG **(a)**, and distinct TG species **(b)**, in resting (M0) macrophages and macrophages stimulated with IFNγ + LPS, measured by MS. **(c, d)** Bodipy 493/503 fluorescence in macrophages stimulated with IFNγ + LPS, with or without DGAT1i, visualized by microscopy **(c)** or measured by flow cytometry **(d)**. **(e)** C16 Bodipy uptake in M0, or macrophages stimulated with IFNγ + LPS, with or without DGATi, measured by flow cytometry. **(f)** Effect of DGAT1i on DG levels in M0 or macrophages stimulated with IFNγ + LPS, measured by MS. **(g)** Effect of Dgat1-shRNA (*shDgat1*) or empty vector (EV) on TG species in inflammatory macrophages, measured by MS. **(h)** Bodipy 493/503 fluorescence in macrophages transduced with sh*Dgatl* or EV and treated with IFNγ + LPS, measured by flow cytometry. **(i, j)** Pro IL-1β positive macrophages stimulated with IFNγ + LPS, plus/minus DGATi (i) or *shDgat1* **(j)**, measured by flow cytometry. **(k, l)** Effect of loss of DGAT1 function on IL-6 production by inflammatory macrophages, measured by ELISA. **(m)** Effect of DGAT1i on phagocytosis of S. *aureus* (PE-labelled), measured by flow cytometry. **(n)** C57BL/6 mice were injected with DGAT1i, or carrier, 30 min before receiving 8mg/kg of i.p. LPS or PBS. Body temperature was monitored every 3 h. Drop in body temperature was calculated as percentage (%) in relation to the initial body temperature. **(o, p)** Bodipy 493/503 fluorescence in large (F4/80+TIM4+) (o), and small (F4/80+TIM4^neg^) **(p)**, peritoneal macrophages from the mice mice shown in n, recovered at 10 h post injection of LPS or PBS. Measured by flow cytometry. **(q)** Serum IL-1β from LPS and LPS + DGAT1i injected mice at 2 h post injection, as measured by ELISA. **(r)** Serum IL-6 from LPS and LPS + DGAT1i injected mice at 10 h post injection, as measured by ELISA. For **a-g** and **k-m**, assays were at 18 h post activation; **i,j** at 6 h. Error bars are mean ± SEM (*p < 0.05, **p < 0.01, ***p < 0.001 and ****p < 0.0001 compared with control or untreated, or as indicated). Representative of two (a-b, e-h, n-r) or three (c-d, i-m) experiments.

We found that inhibiting DGAT1 also had significant effects on inflammation in vivo. T863-treatment of mice with LPS-induced systemic inflammation significantly diminished disease severity, which was measured as the hypothermic response to i.p. LPS injection (Fig. 2n). Mice that had received the drug were active and appeared unaffected by LPS injection at the time of sacrifice (10 h post injection), whereas those that had received carrier alone exhibited sickness signs such as hunching, immobility and piloerection. Post mortem analysis of peritoneal macrophages revealed lower bodipy 493/503 staining in resident (F4/80+ TIM4+) and recruited (F4/80+ TIM4^neg^) macrophages (Figs. 2o, p), indicative of lower LD formation. Moreover, T863-treatment resulted in significantly decreased serum IL-1β (Fig. 2q) and IL-6 (Fig. 2r). Taken together, these results indicate that the synthesis of TGs and their storage in LDs supports inflammatory macrophage activation.

Macrophage activation driven by IFNg plus LPS is linked to increased aerobic glycolysis and the diversion of the TCA cycle intermediates citrate/aconitate to the production of itaconate^6,7^. We reasoned that reduced inflammatory capacity associated with loss of DGAT1 function might be linked to alterations in one or more of these parameters. However, we found that inhibition of TG synthesis had no significant effect on glucose carbon incorporation into lactate (a downstream product of aerobic glycolysis) (Fig. 3a) or citrate (Fig. 3b) or on extracellular acidification rate (ECAR, a measure of released lactate) (Figs. 3c, S3a) or itaconate production (Figs. 3d, S3b) in DGAT1i-treated or *Dgat1*-shRNA-transduced LPS/IFNγ-stimulated cells. We also assessed mitochondrial function. We found that inner membrane potential (a mark of the proton gradient across the inner membrane) (Fig. 3e) and mitochondrial mass (Fig. 3f) were equivalent or greater, respectively, in cells that lacked DGAT1 function, and that mitochondrial reactive oxygen species (ROS) were increased (Figs. 3g, S3c). Consistent with diminished respiration in inflammatory macrophages^17^, ultrastructural analyses revealed that mitochondrial cristae in these cells were looser than in resting cells^22,23^, but this parameter was not obviously affected by DGAT1 inhibition (Fig. 3h; data not shown). As expected, LDs were evident in inflammatory macrophages to a greater extent than in resting macrophages (Fig. S3d), and these organelles were less frequent in cells treated with DGAT1i (Figs. 3h, S3d).

**Figure 3.**
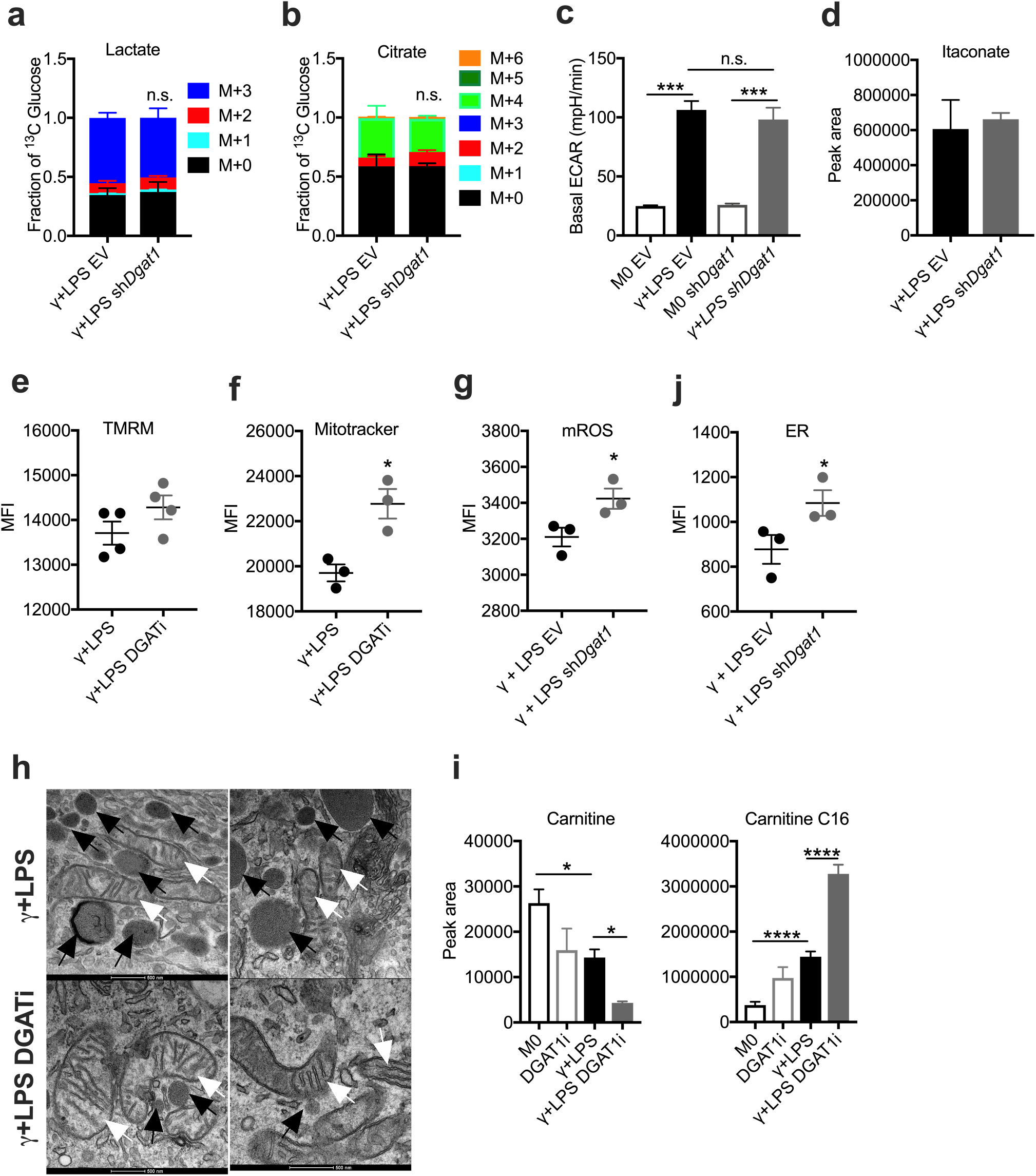
Lack of effect of loss of function of DGAT1 on core metabolism in inflammatory macrophages. **(a, b)** Fraction contribution of glucose carbons into lactate **(a)** and citrate **(b)**, in inflammatory macrophages transduced with EV or sh*Dgat1*, as measured by MS. Macrophages were stimulated with IFNγ + LPS for 18 h and cultured in ^13^C-glucose for the entire time. (c) Basal ECAR of macrophages transduced with EV or sh*Dgatl*, and cultured with IFNγ + LPS or without (M0) for 18 h, measured by extracellular flux analysis. (d) Total pool of itaconate in macrophages transduced with EV or sh*Dgat1* and cultured with IFNγ + LPS or without (M0) for 18 h, as measured by MS. **(e-g)** Effects of DGAT1 loss of function on mitochondrial membrane potential **(e)**, mitochondrial mass **(f)** and mitochondrial reactive oxygen species **(g)**, in macrophages that were stimulated with IFNγ + LPS for 18 h, measured using specific probes and flow cytometry. **(h)** Electron microscopy of macrophages at 18 h post stimulation with IFNγ + LPS without or with DGAT1i. Black arrows, LDs; white arrows, mitochondria. **(i)** Carnitine and C16-acylcarnitine in resting (M0) or IFNγ + LPS-stimulated macrophages, cultured with or without DGAT1i for 18 h. Measured by MS. **(j)** Endoplasmic reticulum (ER) mass in macrophages transduced with EV or sh*Dgat1* and stimulated with IFNγ + LPS for 18 h. Measured using a specific probe and flow cytometry. Error bars are mean ± SEM (\**p* < 0.05, \*\**p* < 0.01, \*\*\**p* < 0.001 and \*\*\*\**p* < 0.0001 compared with control or untreated, or as indicated). **(a-j)** representative of tree experiments.

In other cell types, DGAT1 inhibition has been associated with increased ROS and increased long chain acylcarnitine accumulation^24^. Similarly, we found increased long chain acylcarnitine pools (Figs. 3i, S3e), associated with diminished pools of carnitine (Fig. 3i) and correlated with increased mitochondrial ROS (Figs. 3g, S3c) as a result of loss of DGAT1 function in inflammatory macrophages.

TGs are synthesized in the ER, which we have previously reported is significantly expanded in dendritic cells activated by LPS^25^. We found a similar effect in in macrophages following stimulation with LPS plus IFNγ (Figs. 3j, S3f). Whereas in DCs, ER expansion was shown to require fatty acid synthesis, and to support increased cytokine production, here we found that despite causing diminished cytokine production, inhibition of TG synthesis resulted in an additional increase in ER (Figs. 3j, S3f). We speculate that this may reflect the accumulation of intermediates of the TG synthesis pathway in the ER.

As a means to understand underlying changes in inflammatory function we explored the transcriptional signature associated with loss of DGAT1 function. Inhibition of DGAT1 resulted in significant changes in expression of 285 genes (Fig. S4a). Amongst the six most significantly downregulated pathways within this set, three were linked to eicosanoids, with two specifically associated with the regulation of prostaglandin secretion (Fig. 4a). PGE2 is produced by inflammatory macrophages, in which expression of both prostaglandin synthases *Ptges1* and *Ptges2* was elevated compared to in resting macrophages (Fig. S4c). Previous work has localized PGE2 synthesis to LDs^26^, and postulated that eicosanoid production is a major function of these organelles in immune cells^27^. Moreover, recent findings have implicated autocrine effects of PGE2 in the production of IL-1β^28^. Given that loss of DGAT1 function resulted in loss of LDs (Figs. 2g, h), and of IL-1β production, we asked whether it affected PGE2 levels and thereby inflammatory function. We found that PGE2 production was significantly increased in response to inflammatory signals, and consistent with a role for TGs in PGE2 synthesis, suppressed by DGAT1 inhibition (Fig. 4b), and that exogenous PGE2 increased pro IL-1β and IL-6 production by inflammatory macrophages in the presence of DGAT1i (Figs. 4c, d) or *Dgat1*-shRNA (Figs. S4d, e). There was also a proinflammatory effect of exogenous PGE2 in cells in which DGAT1 was functional (Figs. 4c, d; S4d, e), but the PGE2-induced fold increase in cytokine production was significantly greater in cells in which TG synthesis was inhibited (Figs. 4e, S4f). We found that the addition of exogenous PGE2 also reversed the decline in phagocytosis-competent cells associated with loss of DGAT1 function (Fig. 4f). Moreover, PGE2 increased the intrinsic phagocytic capacity of both control and *Dgat1*-shRNA-transduced inflammatory macrophages (Fig. S4g). Thus, our results indicate that inflammatory macrophages require increased TG synthesis and LD formation in order to allow the synthesis of PGE2 which provides a second signal for inflammatory activity.

**Figure 4.**
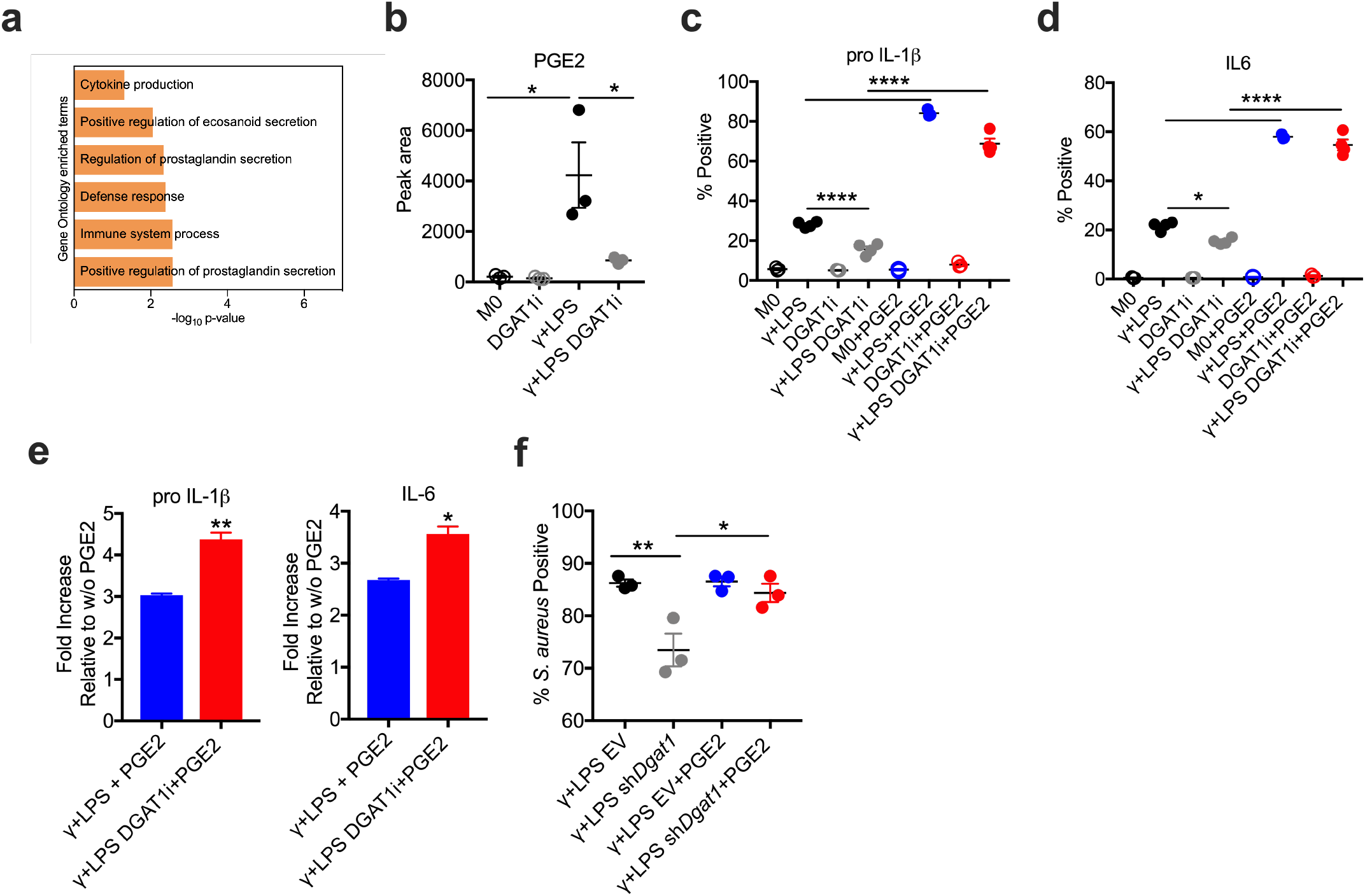
Reduced PGE2 production downstream of inhibited TG synthesis impacts inflammatory macrophage function. **(a)** Down regulated enriched pathways in macrophages stimulated with IFNγ + LPS in the presence or absence of DGAT1i for 18 h. Assessed using RNAseq data. **(b)** Total pool of prostaglandin E2 (PGE2) in resting (M0) or IFNγ + LPS-stimulated macrophages, treated with or without DGAT1i for 18 h. Measured by MS. **(c, d)** pro IL-1β **(c)** or IL-6 **(d)** positive macrophages (percentage cytokine-positive F4/80+ cells, measured by flow cytometry), 6 h after stimulation with IFNγ + LPS with or without DGATi and/or exognenous PGE2, as shown. PGE2 was added 1 hour after IFNγ + LPS or IFNγ + LPS + DGATi, and cells were collected 5 h later. **(e)** Fold increases in pro IL-1β and IL-6 production (measured as percentage cytokine-positive F4/80+ cells) induced by exogenous PGE2 in macrophages stimulated with IFNγ + LPS in the presence or absencer of DGAT1i. **(f)** Effect of PGE2 on the ability of EV or sh*Dgat1*-transduced macrophages to phagocytose S. *aureus* (PE-labelled) after simulation with IFNγ + LPS, measured by flow cytometry. Data show percentage of F4/80+ cells positive for *S.aureus*. Error bars are mean ± SEM (\**p* < 0.05, \*\**p* < 0.01, \*\*\**p* < 0.001 and \*\*\*\**p* < 0.0001 compared with control or untreated, or as indicated). (a) representative of one experiment, (b-f) representative of two experiments.

A surprising finding from the current studies was that inflammatory macrophages have increased pools of long chain acylcarnitines. We reason that this reflects increased substrate levels due to increased FA uptake and synthesis, and subsequent acylcarnitine accumulation due to mitochondrial dysfunction with the cessation of FAO. The functional significance of acylcarnitine accumulation is unknown, but given that long chain acylcarnitines increased further when TG synthesis was inhibited, it may serve as an alternative pathway to LD development (Fig. 1h) for sequestering FA. Increased acylcarnitine accumulation following DGAT1 inhibition has been reported previously in other cell types^29,30^. However, in MEFs, accumulated acylcarnitines due to DGAT1 inhibition were reported to themselves cause mitochondrial dysfunction^29^. Export of long chain acylcarnitines from mitochondria and from cells has been reported^31^, raising the possibility that accumulated acylcarnitines may be released to serve specific extracellular functions, including roles in sterile inflammation^32^ associated with macrophage activation. The dissociation of increased *Cpt1a* expression from FAO is important, since Cpt1a/Cpt-dependent FAO generally works to counteract LD development in macrophages^33^, and enforced expression of a permanently active Cpt1a mutant in a macrophage cell line (RAW) increased FAO and subsequently prevented LD development and inhibited cytokine production in response to an inflammatory signal^34^.

Altered mitochondrial function, in which FAO and OXPHOS are inhibited and the TCA cycle is fragmented to support the use of citrate and aconitate for fatty acid and itaconate production respectively, are core metabolic features of inflammatory macrophage activation^36,7,35,36^. These events, alongside increased FA uptake and increased DGAT expression, create an environment in which TG synthesis is promoted. The finding that LDs, which store TGs, play an important role in maximizing inflammatory macrophage function raises the possibility that LD development is a major objective of metabolic rewiring in these cells. In this scenario, coordinated increases in LD development and *Ptges1* and *Ptges2* expression enable the framework for enhanced PGE2 synthesis, which provides a strong positive feedback signal for the production of pro-IL-1β^28^. IL-1β is considered the gatekeeper of inflammation in both infectious and sterile settings, since it is able to broadly induce the production of other proinflammatory mediators such as TNF, GM-CSF, G-CSF, IL-6, IL-8, NO, as well PGE2 and additional IL-1β^37^. While events that we have shown here to be linked to each other through LDs, such as IL-1β and PGE2 production, have been linked to protective mechanisms in microbial infections^38,39^, there are also clear examples in which exacerbated production of these mediators is linked to the development of severe diseases such as cancer, neurodegenerative diseases, atherosclerotic disease and rheumatologic disorders^40–43^. Our findings raise the possibility of targeting TG synthesis for therapeutic purposes in these settings.

## Acknowledgments

We thank members of the Pearce laboratories, the Electron Microscopy Laboratory at the University of Padova, and the FACS, Imaging, and Deep Sequencing facilities at the Max Planck Institute of Immunobiology and Epigenetics for technical support. This work was funded by by NIH (AI110481 to EJP) and the Max Planck Society. AC is supported by the CAPES/Alexander von Humboldt Fellowship Foundation, MC is supported by the Alexander von Humboldt Fellowship Foundation, MM is supported by Japan Society for the Promotion of Science and LBM is supported by the Sao Paulo research foundation-FAPESP.

## Author Contributions

AC and EJP designed research studies. AC, LBM, NVTB, MC, AMC, FH, MM, GC, RIKG and JB conducted experiments. JEH developed the lipid methodology. DES and NR performed bioinformatics analysis. ELP and EJP provided supervision and funding. AC and EJP wrote the manuscript.

## Methods

### Mice and *in vivo* experiments

C57BL/6 mice (RRID: IMSR_JAX:000664) were from The Jackson Laboratory, and were maintained in specific pathogen-free conditions under protocols approved by the animal care committee of the Regierungspräsidium Freiburg, Germany, in compliance with all relevant ethical regulations. Animals were 6–8 weeks old when used. They were euthanized by carbon dioxide asphyxiation followed by cervical dislocation, and bone marrow, blood and peritoneal lavage were harvested post mortem. For *in vivo* studies, mice were injected i.p. with 8 mg kg^−1^ LPS (Sigma), and body temperature was monitored for the duration of the experiment using an infrared thermometer (Bioseb). Mice were administered T863 (5 mg kg^−1^, i.p.; Sigma) or an equivalent volume of solvent (dimethylsulfoxide, control) 30 mincame prior to LPS injection. Serum and peritoneal lavage were collected at 2 h or 10 h post LPS injection.

### Primary cell cultures

Bone marrow cells were grown in complete medium (RPMI-1640 medium containing 10 mM glucose, 2 mM L-glutamine, 100 U ml^−1^ penicillin-streptomycin and 10% FCS) with 20 ng ml^−1^ murine macrophage colony-stimulating factor 1 (CSF-1; Peprotech) for 7 days, and supplemented with CSF-1 on days 3 and 5. On day 7 macrophages were harvested and then maintained in 20 ng ml^−1^ CSF-1 for subsequent experiments in which they were either maintained in medium without any further additions (M0), or stimulated with with 50 ng ml^−1^ IFN-γ (R&D systems) and 20 ng ml^−1^ LPS (M(γ+LPS)), or 20 ng ml^−1^ IL-4 (Peprotech; MI(L-4)), for 18 h. In some experiments, cells were treated with 50 μM T863 (Sigma) and/or 10 μM PGE2 (Sigma) throughout the 18 h period.

### Lipidomics

The protocol for lipid extraction was adapted from Matyash *et al*.^44^. Frozen cell pellets (5×10^5^ cells) were resuspended in ice cold PBS and transferred to glass tubes before the addition of methanol and methyl tert-butyl ether. The tubes were then shaken for 1 h at 4°C. Water was added to separate the phases before centrifugation at 1,000 x g for 10 min. The upper organic phase was collected and dried in a Genevac EZ2 speed vac. Samples were resuspended in 2:1:1 isopropanol:acetonitrile:water prior to analysis. LC-MS was carried out using an Agilent Zorbax Eclipse Plus C18 column using an Agilent 1290 Infinity II UHPLC inline with an Agilent 6495 Triple Quad QQQ-MS. Lipids were identified by fragmentation and retention time, and were quantified using Agilent Mass Hunter software.

### Metabolite quantification

Targeted metabolite quatification was carried out using the same LC-MS machine that was used for lipidomics. Samples were extracted in 50:30:20 methanol:acetonitrile:H2O pre-cooled at −80 °C. Multiple reaction monitoring (MRM) settings were optimized for all metabolites using pure compounds. LC separation was on a Waters CSH-C18 column (100 x 2.1 mm, 1.7 μm particles) using a binary solvent gradient of 100 % buffer A (0.1 % formic acid in water) to 97 % buffer B (50:50 methanol:acetonitrile). Flow rate was 400 μL/min, autosampler temperature was 4 °C, and injection volume was 3 μL. Data processing was performed by an in-house R script. Peak areas were normalized to a fully ^13^C labelled yeast extract (ISOtopic Solutions, Vienna).

### Palmitate tracing

Cells (3×10^5^) were cultured in RPMI 1640, supplemented with 10 mM ^13^C-palmitate for 18 h, after which they were rinsed with cold 0.9% NaCl and extracted using 0.2 mL of 80% MeOH on dry ice, and dried using a SpeedVac. Label tracing was carried out using an Agilent 1290 Infinity II UHPLC inline with a Bruker impact II QTOF-MS operated in full scan (MS1) mode. LC parameters were identical to those used for targeted metabolite quantification. Data processing including correction for natural isotope abundance was performed by an in-house R script. Metabolite peaks were identified based on exact mass and matching of retention time to a pure standard.

### Glucose and glycerol tracing

Cells (3×10^5^) were cultured in RPMI 1640, supplemented with 10mM ^13^C-glucose for 18 h. Separately, media was supplemented with 10mM ^13^C-glycerol for 30 min prior to collection of samples at 18 h post stimulation. Cells were rinsed with cold 0.9% NaCl and extracted using 0.2 mL of 80% MeOH on dry ice. 10 nM norvaline (internal standard) was added. Following mixing and centrifugation, the supernatant was collected and dried using a SpeedVac. Dried extracts were analyzed using gas chromatography-mass spectrometry (GC-MS) (Agilent 5977). Correction for natural isotope abundance and calculation of fractional contribution was performed as described elsewhere^45^.

### Lentiviral and retroviral production and cell transduction

HEK293T cells were transfected using Lipofectamine 3000 (Thermo Fisher Scientific) with lentiviral packaging vectors pCAG-eco and psPAX.2 plus empty pLKO.1 control (EV) with a puromycin selection cassette (all obtained from Addgene) or a shRNA containing pLKO.1 targeting DGAT1 (GE Dharmacon CAT# RMM4534-EG13350 - TRCN0000124791). Virus was collected from the supernatant of the cells. Bone marrow cultures were transduced in the presence of polybrene (8 mg ml–1) on day 2 of culture. A 48 h, selection of transduced cells was performed with 6 μg ml^−1^ puromycin (Sigma).

### RNA sequencing

Total RNA was extracted with the RNAqueous-Micro Total RNA Isolation kit (Thermo Fisher Scientific) and quantified using Qubit 2.0 (Thermo Fisher Scientific) according to the manufacturer’s instructions. Libraries were prepared using the TruSeq stranded mRNA kit (Illumina) and sequenced in a HISeq 3000 (Illumina) by the Deep Sequencing Facility at the Max Planck Institute for Immunobiology and Epigenetics. Sequenced libraries were processed with a pipeline optimized by the Bioinformatics core at the Max Planck Institute for Immunobiology and Epigenetics^46^. Raw mapped reads were processed in R (Lucent Technologies) with DESeq2^47^ to determine differentially expressed genes and generate normalized read counts to visualize as heatmaps using Morpheus (Broad Institute).

### Flow cytometry

Cells were incubated in 5 μg ml^−1^ anti-CD16/32 (Biozol), stained with Live Dead Fixable Blue (Life Technologies), and then surface stained with a fluorochrome-conjugated monoclonal antibody to F4/80 (eBioscience or Biozol, clone BM8), prior to fixation and permeabilization using BD Cytofix/Cytoperm kit (BD Biosciences) and staining with monoclonal antibodies to IL-6 (clone MP5-20F3) and pro IL-1β (clone NJTEN3). Resting (M0) macrophages and macrophages stimulated with IFNγ + LPS, with or without DGAT1i 1 h priori to the addition of Brefeldin A (1:1000) were collected 5 h later. For ICS in cells treated with PGE2: cells were stimulated with IFNγ + LPS 1 h prior to tha addition of PGE2 (10μM) and 1 h later Brefeldin A was added to the culture and ICS was performed after 5 h of subsequent culture. For mitochondrial superoxide staining and ER staining, macrophages were incubated with 5 μM MitoSOX (Thermo Fisher Scientific) or 1 μM ER tracker green (Thermo Fisher Scientific) in complete media without FCS for 10 min for MitoSOX staining and in complete media with FCS for 30 min for ER staining. pHrodo^™^ Red S. *aureus* Bioparticles^™^ (ThermoFisher) was used for phagocytosis assays according to the manufacturer’s instructions. BODIPY^™^ 493/503 (ThermoFisher) and BODIPY^®^ FL C16 (ThermoFisher) were used for neutral lipid staining and fatty acid uptake assays according to the manufacturer’s instructions. Data were acquired by flow cytometry on an LSRII or LSR Fortessa (BD Biosciences) and analyzed with FlowJo v.10.1 (Tree Star).

### Confocal microscopy

2 × 10^5^ macrophages were plated on a 10 mm cover slip in a 24-well plate. Cells were subsequently cultured with or without γ+LPS for 18 h, followed by staining with Bodipy 493/503 for 1 h, fixation in 2% PFA for 20 min and washing with PBS. Slips were mounted with Vectashield with DAPI and sealed with nail polish before image acquisition using a Zeiss LSM 880 with Aryscan equipped with a 63× objective. Confocal images were analyzed and merged using ImageJ software.

### Electron Microscopy

5 × 10^5^ macrophages were fixed in 2.5% glutaraldehyde (Sigma) in 100 mM sodium cocodylate (Sigma) and washed in cocodylate buffer. Following this step, samples were processed and imaged at the Electron Microscopy Laboratory at the University of Padova. LDs were analysed over 50 independent images.

### ELISA

Concentrations of IL-1β and IL-6 in cell culture supernatants or blood serum were measured using cytokine-specific ELISAs (BioLegend ELISA Max kits), according to the manufacturer’s instructions. Absorbance was measured using a TriStar plate reader (Berthold Technologies). Standard curves were analyzed in Prism using second-order polynomial interpolation.

### ECAR measurements

ECAR measurements were made with an XF-96 Extracellular Flux Analyzer (Seahorse Bioscience). 1 × 10^5^ BMDMs, were plated into each well of Seahorse X96 cell culture microplate and preincubated at 37 °C for a minimum of 45 min in the absence of CO2 in unbuffered RPMI with 1 mM pyruvate, 2 mM L-glutamine and 25 mM glucose, with pH adjusted to 7.4. ECAR was measured under basal conditions. Results were collected with Wave software version 2.4 (Agilent).

### Gene expression analysis by quantitative PCR with reverse transcription

RNA was isolated using the RNeasy kit (Qiagen) and single-strand cDNA was synthesized using the High Capacity cDNA Reverse Transcription Kit (Applied Biosystems). DGAT1 RT-PCR was performed with Taqman primerds using an Applied Biosystems 7000 sequence detection system. The expression levels of mRNA were normalized to the expression of β-actin.

### Statistical Analysis

Statistical analysis was performed using prism 6 software (Graph pad) and results are represented as mean ± SEM. Comparisons for two groups were calculated using unpaired two-tailed Student’s t tests, comparisons of more than two groups were calculated using one-way ANOVA with Bonferroni’s multiple comparison tests. We observed normal distribution and no difference in variance between groups in individual comparisons. Statistical significance: *p<0.05; ** p < 0.01; *** p < 0.001; **** p < 0.0001. Further details on statistical analysis are listed in the figure legends.

## Supplemental Figures Legends

**Supplemental Figure 1.**
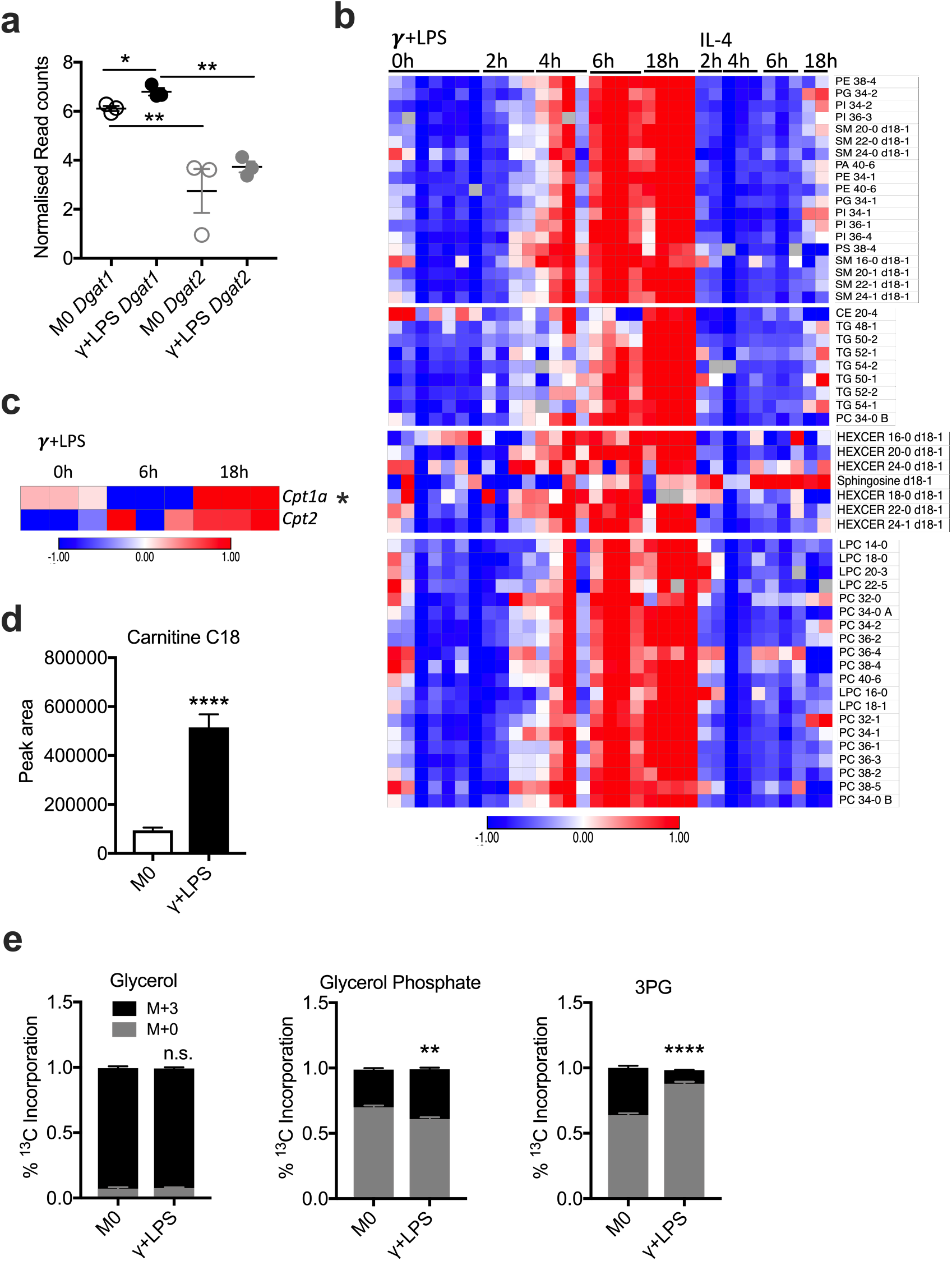
Increased *Dgat1* gene expression, lipid content and incorporation of glycerol into TG synthesis in inflammatory macrophages. **(a)** *Dgat1* and *Dgat2* gene expression in macrophages treated without (M0) or with IFNγ + LPS for 18 h. Data from RNAseq. **(b)** Time-dependent accumulation of different lipid species in macrophages after 0, 2, 4, 6 and 18 h of IFNγ + LPS or IL-4 stimulation. Measured by MS. **(c)** *Cpt1a* and *Cpt2* expression in resting macrophages (0 h) and after 6 h or 18 h of stimulation with IFNγ + LPS (γ + LPS). Data from RNA-seq. **(d)** C18 acylcarnitine measured using MS in M0 macrophages and macrophages stimulated with IFNγ + LPS, for 18 h, **(e)** Fractional contribution of ^13^C Glycerol during a 30 min pulse of labeling resting macrophages (M0) or macrophages stimulated with IFNγ + LPS for 18 h. Error bars are mean ± SEM (\**p* < 0.05, \*\**p* < 0.01, \*\*\**p* < 0.001 and \*\*\*\**p* < 0.0001 compared with control or untreated, or as indicated). (a-e) representative of two experiments.

**Supplemental Figure 2.**
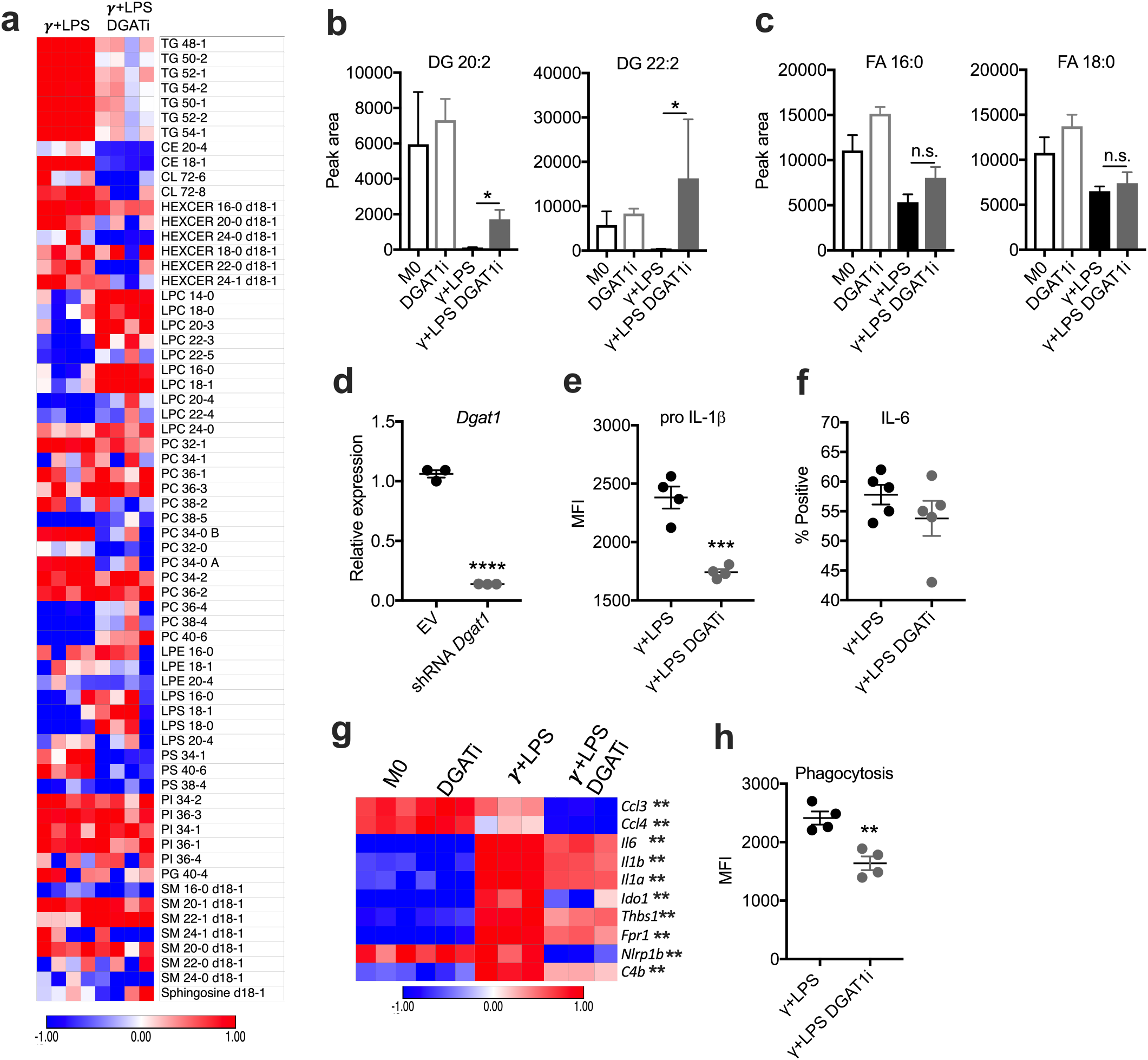
Changes in lipid content of inflammatory macrophages as a result of loss of DGAT1 function. **(a)** Changes in the lipidome of inflammatory macrophages due to inhibition of DGAT1. Measured by MS at 18 h after addition of IFNγ + LPS without or with DGAT1i. **(b, c)** Effect of DGAT1i on DG (DG 20:2 and 22:2) **(b)**, and FA (FA 16:0 and 18:0) **(c)**, in macrophages over 18 h of culture with IFNγ + LPS. Measured by MS. **(d)** Validation of ability of sh*Dgat1* to suppress *Dgat1* gene expression after 3 days of puromycin selection. **(e, f)** pro IL-1β production, measured as MFI by flow cytometry **(e)**, or IL-6 production (percentage cytokine-positive F4/80+ cells, measured by flow cytometry) **(f)**, 6 h after stimulation with IFNγ + LPS with or without DGATi. **(g)** Down regulated genes related to cytokine production and immune system processes in resting (M0) or IFNγ + LPS-stimulated macrophagescultured with or without DGAT1i for 18 h. **(h)** Effects of DGAT1i on the ability of inflammatory macrophages to phagocytose S. *aureus* (PE-labelled), measured by flow cytometry. Data show MFI for *S.aureus* fluorescence in F4/80+ cells. Macrophages were stimulated with IFNγ + LPS for 18 h with or without DAGT1i and then incubated with PE-labeled S.*aureus* (PE) for 30 min. Error bars are mean ± SEM (\**p* < 0.05, \*\**p* < 0.01, \*\*\**p* < 0.001 and \*\*\*\**p* < 0.0001 compared with control or untreated, or as indicated). (a-d) representative of two experiments, (g) representative of one experiment and (e, f, h) representative of tree experiments.

**Supplemental Figure 3.**
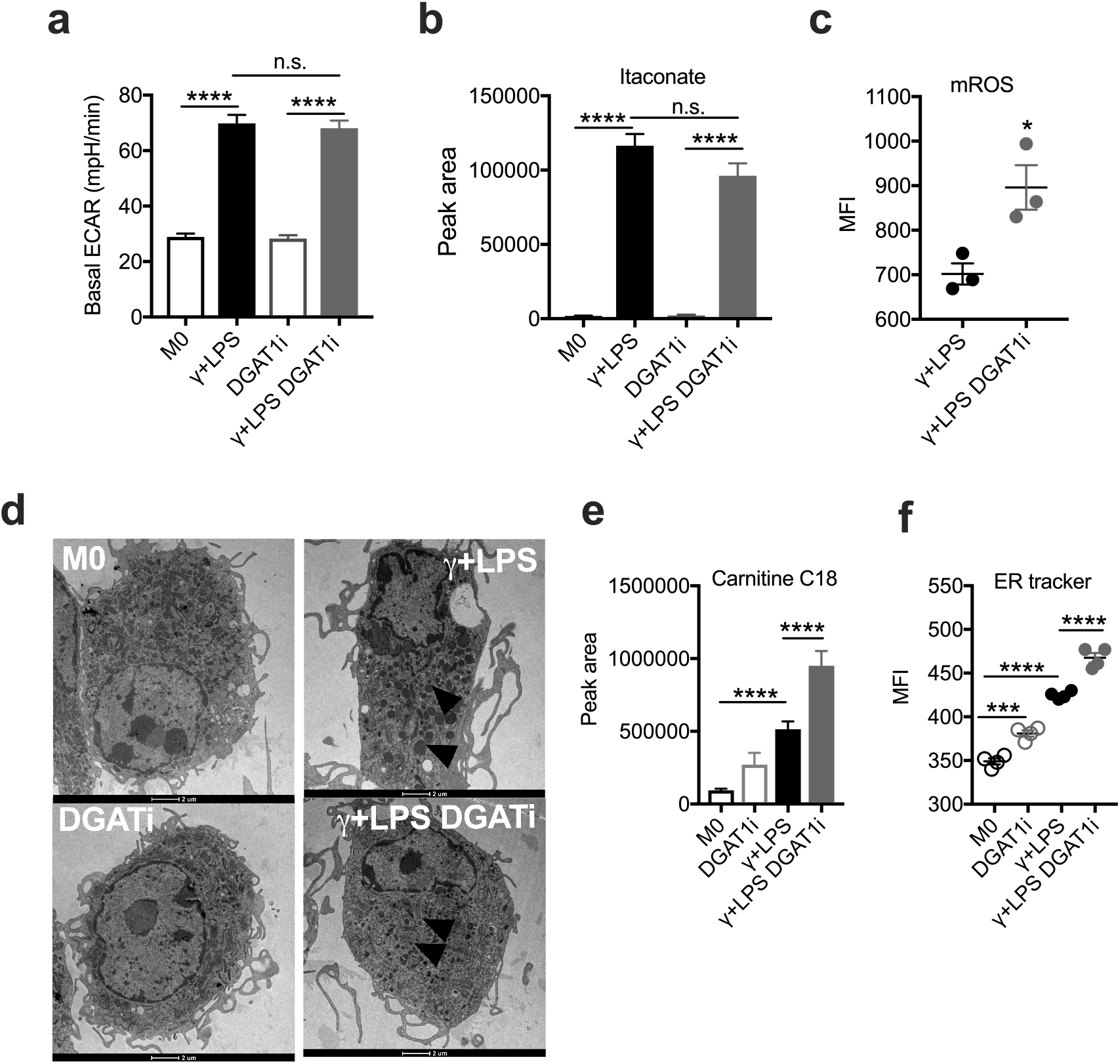
Loss of function of DGAT1 does not change overall cell metabolism in inflammatory macrophages. **(a)** Basal ECAR of macrophages cultured with IFNγ + LPS, or without (M0), with or without DGAT1i for 18 h. Measured by extracellular flux analysis. **(b)** Total pool of itaconate in macrophages cultured with IFNγ + LPS, or without (M0), with or without DGAT1i for 18 h, as measured by MS. **(c)** Effects of DGATi treatment on mitochondrial reactive oxygen species. Macrophages were cultured with IFNγ + LPS for 18 h, after which mitochondrial ROS was measured, using flow cytometry, with a specific probe. **(d)** Low magnification electron micrographs of resting (M0) macrophages or IFNγ + LPS-stimulated macrophages, cultured with or without DGAT1i for 18 h, to show changes in LDs associated with treatments. **(e)** Carnitine and C18-acylcarnitine in resting (M0) macrophages or IFNγ + LPS-stimulated macrophages, cultured with or without DGAT1i for 18 h. (f) Endoplasmic reticulum (ER) mass in macrophages cultured with IFNγ + LPS, or without (M0), with or without DGAT1i for 18h. Measured using a specific probe and flow cytometry. Error bars are mean ± SEM (\**p* < 0.05, \*\**p* < 0.01, \*\*\**p* < 0.001 and \*\*\*\**p* < 0.0001 compared with control or untreated, or as indicated). (a-f) representative of tree experiments.

**Supplemental Figure 4.:**
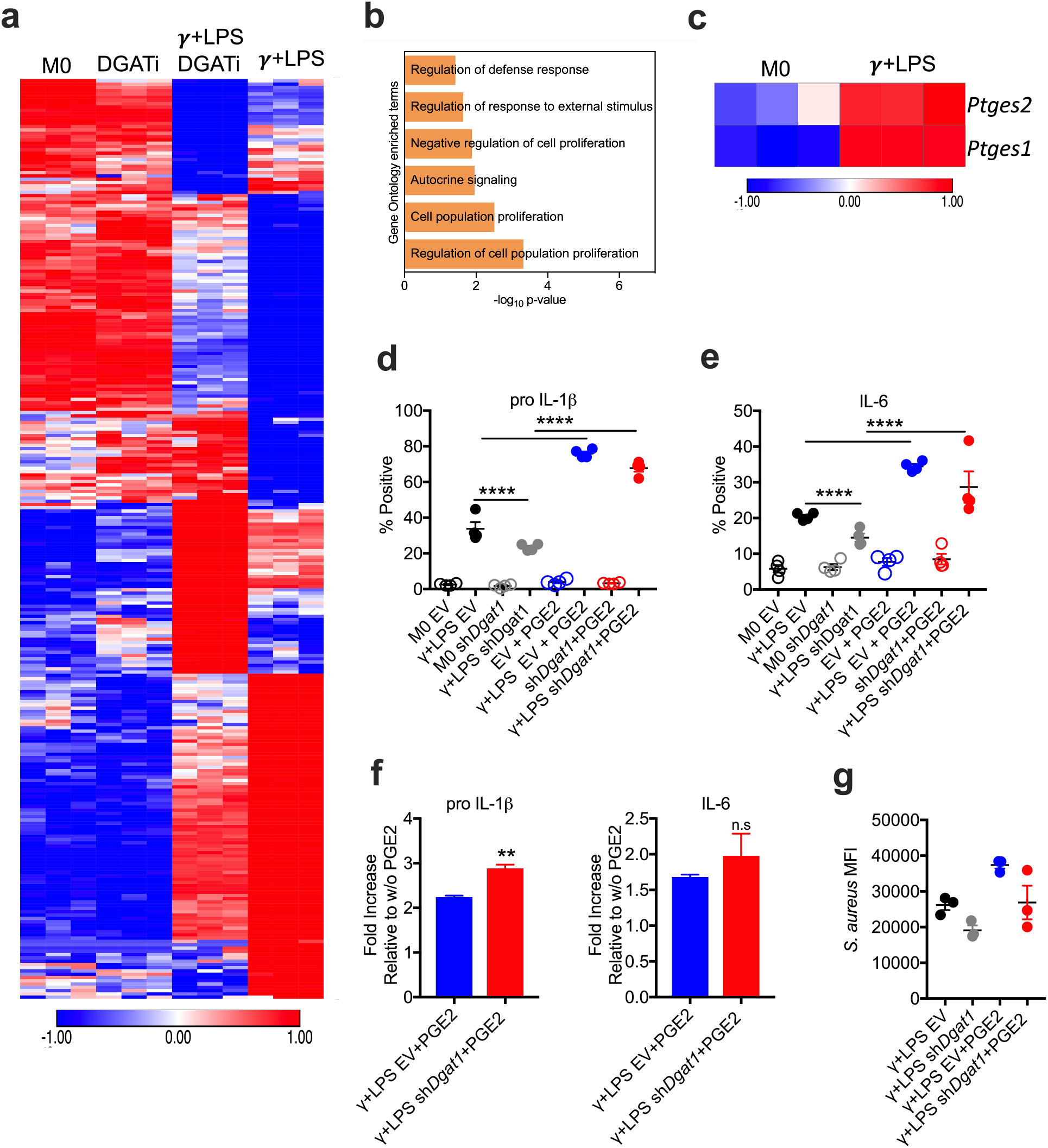
Down- and up-regulated genes and pathways in macrophages with loss of function of *DGAT1*. **(a)** Significantly regulated (p<0.01) genes in macrophages stimulated without (M0) or with IFNγ + LPS for 18 h in the presence or absence of DGAT1i. 136 genes down-regulated and 149 genes up-regulated; data from RNA-seq. **(b)** Up-regulated pathways in macrophages stimulated with IFNγ + LPS versus macrophages stimulated γ + LPS and treated with DGAT1i (IFNγ + LPS DGAT1i) for 18 h. **(c)** *Ptges1* and *Ptges2* expression in resting (M0) macrophages and macrophages treated with IFNγ + LPS for 18 h. Data from RNA-seq. **(d, e)** Macrophages were transduced with empty vector or sh*Dgat1* and cultured with or without IFNγ + LPS, with or without exogenous PGE2 (added 1 h after IFNγ + LPS) for 5 h. Pro IL-1β **(d)** or IL-6 **(e)** positive macrophages (percentage cytokine-positive F4/80+ cells, (measured by flow cytometry), are shown. **(f)** Fold increases in pro IL-1β and IL-6 production (measured as percentage cytokine-positive F4/80+ cells) induced by exogenous PGE2 in macrophages treated as shown. **(g)** Effect of PGE2 on the ability of EV- or sh*Dgat1*-transduced macrophages to phagocytose S. *aureus* (PE-labelled) after simulation with IFNγ + LPS, *as* measured by flow cytometry. Data show MFI values of S.aureus-associated fluorescence in F4/80+ cells. Error bars are mean ± SEM (\**p* < 0.05, \*\**p* < 0.01, \*\*\**p* < 0.001 and \*\*\*\**p* < 0.0001 compared with control or untreated, or as indicated). (a-b) representative of one experiment, (c-g) representative of two experiments.

## References

1. Guerrini V, Gennaro ML. Foam Cells: One Size Doesn’t Fit All. Trends Immunol. 2019.

2. Thiam AR, Farese RV, Walther TC. The biophysics and cell biology of lipid droplets. Nat Rev Mol Cell Biol. 2013;14(12):775–786.

3. Huang YL, Morales-Rosado J, Ray J, et al. Toll-like receptor agonists promote prolonged triglyceride storage in macrophages. J Biol Chem. 2014;289(5):3001–3012.

4. Huang SC, Everts B, Ivanova Y, et al. Cell-intrinsic lysosomal lipolysis is essential for alternative activation of macrophages. Nat Immunol. 2014;15(9):846–855.

5. Dinarello CA. Interleukin-1 in the pathogenesis and treatment of inflammatory diseases. Blood. 2011;117(14):3720–3732.

6. O’Neill LA, Pearce EJ. Immunometabolism governs dendritic cell and macrophage function. J Exp Med. 2016;213(1):15–23.

7. Jha AK, Huang SC, Sergushichev A, et al. Network integration of parallel metabolic and transcriptional data reveals metabolic modules that regulate macrophage polarization. Immunity. 2015;42(3):419–430.

8. Dennis EA, Deems RA, Harkewicz R, et al. A mouse macrophage lipidome. J Biol Chem. 2010;285(51):39976–39985.

9. Funk JL, Feingold KR, Moser AH, Grunfeld C. Lipopolysaccharide stimulation of RAW 264.7 macrophages induces lipid accumulation and foam cell formation. Atherosclerosis. 1993;98(1):67–82.

10. Yen CL, Stone SJ, Koliwad S, Harris C, Farese RV. Thematic review series: glycerolipids. DGAT enzymes and triacylglycerol biosynthesis. J Lipid Res. 2008;49(11):2283–2301.

11. Henne WM, Reese ML, Goodman JM. The assembly of lipid droplets and their roles in challenged cells. EMBO J. 2018;37(12).

12. Chitraju C, Mejhert N, Haas JT, et al. Triglyceride Synthesis by DGAT1 Protects Adipocytes from Lipid-Induced ER Stress during Lipolysis. Cell Metab. 2017;26(2):407–418.e403.

13. Piccolis M, Bond LM, Kampmann M, et al. Probing the Global Cellular Responses to Lipotoxicity Caused by Saturated Fatty Acids. Mol Cell. 2019;74(1):32–44.e38.

14. Li AC, Glass CK. The macrophage foam cell as a target for therapeutic intervention. Nat Med. 2002;8(11):1235–1242.

15. Childs BG, Baker DJ, Wijshake T, Conover CA, Campisi J, van Deursen JM. Senescent intimal foam cells are deleterious at all stages of atherosclerosis. Science. 2016;354(6311):472–477.

16. Kim K, Shim D, Lee JS, et al. Transcriptome Analysis Reveals Nonfoamy Rather Than Foamy Plaque Macrophages Are Proinflammatory in Atherosclerotic Murine Models. Circ Res. 2018;123(10):1127–1142.

17. Cameron AM, Castoldi A, Sanin DE, et al. Inflammatory macrophage dependence on NAD. Nat Immunol. 2019;20(4):420–432.

18. Cao J, Zhou Y, Peng H, et al. Targeting Acyl-CoA:diacylglycerol acyltransferase 1 (DGAT1) with small molecule inhibitors for the treatment of metabolic diseases. J Biol Chem. 2011;286(48):41838–41851.

19. Jaisinghani N, Dawa S, Singh K, et al. Necrosis Driven Triglyceride Synthesis Primes Macrophages for Inflammation During. Front Immunol. 2018;9:1490.

20. Freigang S, Ampenberger F, Weiss A, et al. Fatty acid-induced mitochondrial uncoupling elicits inflammasome-independent IL-1α and sterile vascular inflammation in atherosclerosis. Nat Immunol. 2013;14(10):1045–1053.

21. Chandak PG, Radovic B, Aflaki E, et al. Efficient phagocytosis requires triacylglycerol hydrolysis by adipose triglyceride lipase. J Biol Chem. 2010;285(26):20192–20201.

22. Cogliati S, Frezza C, Soriano ME, et al. Mitochondrial cristae shape determines respiratory chain supercomplexes assembly and respiratory efficiency. Cell. 2013;155(1):160–171.

23. Buck MD, O’Sullivan D, Klein Geltink RI, et al. Mitochondrial Dynamics Controls T Cell Fate through Metabolic Programming. Cell. 2016;166(1):63–76.

24. Liepinsh E, Makrecka-Kuka M, Volska K, et al. Long-chain acylcarnitines determine ischaemia/reperfusion-induced damage in heart mitochondria. Biochem J. 2016;473(9):1191–1202.

25. Everts B, Amiel E, Huang SC, et al. TLR-driven early glycolytic reprogramming via the kinases TBK1-IKKε supports the anabolic demands of dendritic cell activation. Nat Immunol. 2014;15(4):323–332.

26. Accioly MT, Pacheco P, Maya-Monteiro CM, et al. Lipid bodies are reservoirs of cyclooxygenase-2 and sites of prostaglandin-E2 synthesis in colon cancer cells. Cancer Res. 2008;68(6):1732–1740.

27. Melo RC, Weller PF. Lipid droplets in leukocytes: Organelles linked to inflammatory responses. Exp Cell Res. 2016;340(2):193–197.

28. Zasłona Z, Pålsson-McDermott EM, Menon D, et al. The Induction of Pro-IL-1β by Lipopolysaccharide Requires Endogenous Prostaglandin E. J Immunol. 2017;198(9):3558–3564.

29. Nguyen TB, Louie SM, Daniele JR, et al. DGAT1-Dependent Lipid Droplet Biogenesis Protects Mitochondrial Function during Starvation-Induced Autophagy. Dev Cell. 2017;42(1):9–21.e25.

30. Ackerman D, Tumanov S, Qiu B, et al. Triglycerides Promote Lipid Homeostasis during Hypoxic Stress by Balancing Fatty Acid Saturation. Cell Rep. 2018;24(10):2596–2605.e2595.

31. Violante S, Ijlst L, Te Brinke H, et al. Carnitine palmitoyltransferase 2 and carnitine/acylcarnitine translocase are involved in the mitochondrial synthesis and export of acylcarnitines. FASEB J. 2013;27(5):2039–2044.

32. Rutkowsky JM, Knotts TA, Ono-Moore KD, et al. Acylcarnitines activate proinflammatory signaling pathways. Am J Physiol Endocrinol Metab. 2014;306(12):E1378–1387.

33. Nomura M, Liu J, Yu ZX, et al. Macrophage fatty acid oxidation inhibits atherosclerosis progression. J Mol Cell Cardiol. 2019;127:270–276.

34. Malandrino MI, Fucho R, Weber M, et al. Enhanced fatty acid oxidation in adipocytes and macrophages reduces lipid-induced triglyceride accumulation and inflammation. Am J Physiol Endocrinol Metab. 2015;308(9):E756–769.

35. O’Neill LAJ, Artyomov MN. Itaconate: the poster child of metabolic reprogramming in macrophage function. Nat Rev Immunol. 2019;19(5):273–281.

36. Williams NC, O’Neill LAJ. A Role for the Krebs Cycle Intermediate Citrate in Metabolic Reprogramming in Innate Immunity and Inflammation. Front Immunol. 2018;9:141.

37. Dinarello CA. A clinical perspective of IL-1β as the gatekeeper of inflammation. Eur J Immunol. 2011;41(5):1203–1217.

38. Chen M, Divangahi M, Gan H, et al. Lipid mediators in innate immunity against tuberculosis: opposing roles of PGE2 and LXA4 in the induction of macrophage death. J Exp Med. 2008;205(12):2791–2801.

39. Ridker PM, MacFadyen JG, Thuren T, et al. Effect of interleukin-1β inhibition with canakinumab on incident lung cancer in patients with atherosclerosis: exploratory results from a randomised, double-blind, placebo-controlled trial. Lancet. 2017;390(10105):1833–1842.

40. Ridker PM, Everett BM, Thuren T, et al. Antiinflammatory Therapy with Canakinumab for Atherosclerotic Disease. N Engl J Med. 2017;377(12): 1119–1131.

41. Lachmann HJ, Kone-Paut I, Kuemmerle-Deschner JB, et al. Use of canakinumab in the cryopyrin-associated periodic syndrome. N Engl J Med. 2009;360(23):2416–2425.

42. Leow MK. PTEN haploinsufficiency, obesity, and insulin sensitivity. N Engl J Med. 2012;367(25):2450–2451; author reply 2451.

43. Yagami T, Koma H, Yamamoto Y. Pathophysiological Roles of Cyclooxygenases and Prostaglandins in the Central Nervous System. Mol Neurobiol. 2016;53(7):4754–4771.

44. Matyash V, Liebisch G, Kurzchalia TV, Shevchenko A, Schwudke D. Lipid extraction by methyl-tert-butyl ether for high-throughput lipidomics. J Lipid Res. 2008;49(5):1137–1146.

45. Buescher JM, Antoniewicz MR, Boros LG, et al. A roadmap for interpreting (13)C metabolite labeling patterns from cells. Curr Opin Biotechnol. 2015;34:189–201.

46. Bhardwaj V, Heyne S, Sikora K, et al. snakePipes: facilitating flexible, scalable and integrative epigenomic analysis. Bioinformatics. 2019;35(22):4757–4759.

47. Love MI, Huber W, Anders S. Moderated estimation of fold change and dispersion for RNA-seq data with DESeq2. Genome Biol. 2014;15(12):550.

